# Puromycin-sensitive aminopeptidase acts as an inhibitory auxiliary subunit of volume-regulated anion channels

**DOI:** 10.1101/2025.08.24.671966

**Authors:** Wenqiang Zheng, Tatsuya Hagino, Hao Wang, Henry Yi Cheng, Nicholas Koylass, Kevin Hong Chen, Haobo Wang, Sepehr Mani, Anish Kumar Mondal, Edward C. Twomey, Zhaozhu Qiu

## Abstract

Volume-regulated anion channels (VRACs) are large-pore channels present in nearly all vertebrate cells, playing key roles in cell volume regulation and autocrine/paracrine signaling. Here, we identify the ubiquitously expressed puromycin-sensitive aminopeptidase (PSA) as a binding partner of the obligatory VRAC subunit SWELL1 (also known as LRRC8A) and report the cryo-electron microscopy structure of the SWELL1–PSA complex. Three PSA molecules associate with a single SWELL1 hexamer, coupling adjacent leucine-rich repeat (LRR) domains into local dimers. Functionally, PSA overexpression suppresses VRAC activation, whereas its deletion results in elevated basal channel activity. Notably, PSA’s regulatory role on VRACs is independent of its aminopeptidase activity. Our findings identify PSA as the first auxiliary subunit of VRACs, highlight the role of intracellular LRR domains in allosteric channel gating, and propose a new strategy for modulating VRAC function in diverse physiological contexts, including cGAMP transport and STING signaling.

## INTRODUCTION

Volume-regulated anion channels (VRACs) are activated by cell swelling to facilitate the efflux of chloride ions and organic osmolytes, playing a central role in the cellular response to osmotic stress^1–3^. Beyond their canonical function in cell volume regulation, VRACs mediate the transport of diverse signaling molecules—including 2’3’-cyclic-GMP-AMP (cGAMP), glutamate, GABA, and ATP—as well as drugs such as cisplatin^4–10^. By doing so, VRACs influence a wide array of physiological and pathological processes, including immune responses^4,11^, neuron-glia interactions^5,6^, metabolism^12,13^, and cancer^8^.

SWELL1 (LRRC8A), a member of the leucine-rich repeat-containing family 8 (LRRC8) proteins, is the main pore-forming subunit of VRACs^14,15^, essential for their localization to the plasma membrane. It assembles into hetero-hexameric channel complexes with one or more closely related paralogues (LRRC8B–E)^14,16^. The specific subunit composition determines the channels’ biophysical properties, including conductance and substrate selectivity^9,16^. While many ion channels also stably interact with non-pore-forming auxiliary subunits to modulate their trafficking and gating properties, none have been identified for VRACs. However, the cytoplasmic leucine-rich repeat (LRR) domains of VRACs are well-established scaffolds for protein-protein interactions^17^. Notably, several synthetic nanobodies (sybodies) have been generated to bind distinct regions of the LRR domains and modulate VRAC activity^18^, supporting the possibility that native proteins may similarly regulate the channels through allosteric mechanisms. However, to date, no endogenous VRAC auxiliary subunits have been identified.

To investigate how VRACs are regulated under native conditions, we performed proteomics analyses and identified puromycin-sensitive aminopeptidase (PSA; gene name: *NPEPPS*) as a binding partner of SWELL1. Cryo-electron microscopy (cryo-EM) revealed that a single PSA molecule engages two adjacent leucine-rich repeat (LRR) domains of SWELL1, stabilizing a dimeric configuration. Electrophysiological studies demonstrated that PSA inhibits VRAC activity, and functional assays showed that PSA modulates cGAMP transport through this regulatory mechanism. Notably, a recent study reported that PSA/NPEPPS interacts with VRACs to regulate cisplatin import and modulate cisplatin sensitivity across multiple tumor types^19^, underscoring the broader physiological and therapeutic relevance of this interaction. By establishing PSA as an inhibitory auxiliary subunit of VRACs, our work provides a structural and mechanistic framework for understanding VRAC regulation in both physiology and disease.

## RESULTS

### Identification of PSA as a SWELL1-binding protein

Since SWELL1 is the principal component of VRACs, we overexpressed SWELL1-Flag in HEK293 cells and performed affinity purification using an anti-Flag antibody followed by gel electrophoresis and mass spectrometry (MS) to identify SWELL1-interacting partners. In addition to SWELL1 itself, one of the most prominent binding proteins identified was puromycin-sensitive aminopeptidase (PSA, also known as NPEPPS), a ubiquitously expressed cytosolic M1 family metallopeptidase that cleaves N-terminal amino acids from peptides (**Figure 1A**). Among the VRAC subunits, only LRRC8C was detected, likely due to the low abundance of native LRRC8 proteins.

**Figure 1.**
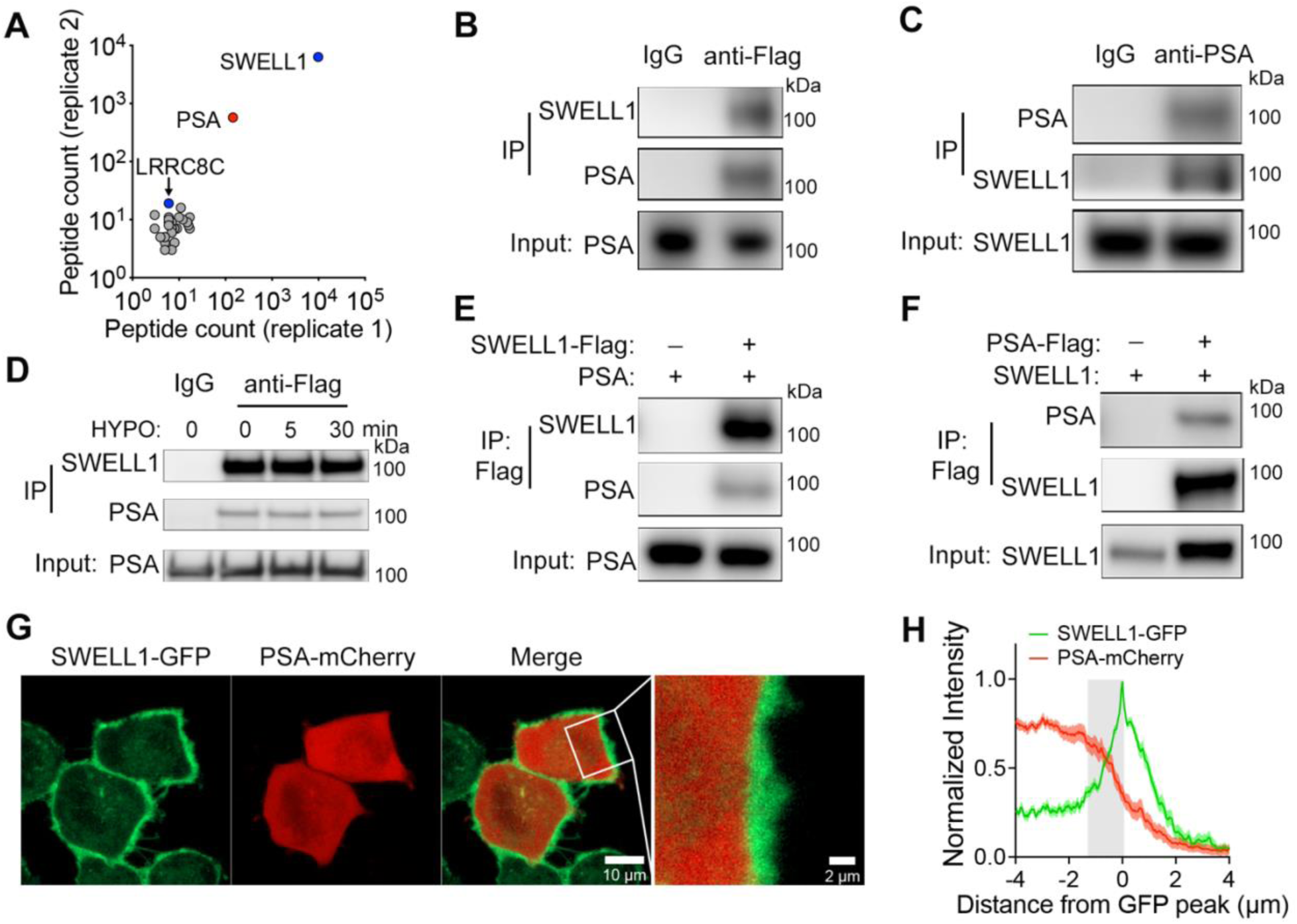
Identification of PSA as a SWELL1-binding partner. (A) Unique peptide counts of common hits (≥3 distinct peptides absent from control samples) identified in both replicates by mass spectrometry of affinity-purified samples from SWELL1-Flag overexpressing cells. Three top hits are annotated (B and C) Immunoprecipitation (IP) followed by immunoblotting showing that anti-Flag antibodies pull down PSA (B) and anti-PSA antibodies pull down SWELL1 (C) in SWELL1-Flag knock-in HeLa cells. IgG was used as a negative control. (D) The co-IP levels of PSA were not changed during cell swelling. SWELL1-Flag knock-in HeLa cells were treated with hypotonic solution (230 mOsm/kg) for 0, 5, or 30 min prior to cell lysis. (E and F) IP followed by immunoblot showing that SWELL1 co-precipitates with PSA (E) and PSA co-precipitates with SWELL1 (F) when SWELL1 and PSA plasmids were co-transfected into *LRRC8⁻/⁻* HEK293 cells. (G) Representative live cell images of SWELL1-GFP knock-in HeLa cells expressing PSA-mCherry fusion protein. (H)Normalized fluorescence intensities (mean ± SEM, n = 24 cells) measured along line scans through the plasma membrane.

To further validate this interaction, we used HeLa cells with a Flag tag inserted into the endogenous *SWELL1* locus^16^, thereby circumventing the lack of a suitable immunoprecipitation (IP)-grade anti-SWELL1 antibody. IP followed by immunoblot confirmed that anti-Flag antibody co-precipitates PSA (**Figure 1B**), and conversely, anti-PSA antibody pulls down SWELL1 (**Figure 1C**). These results indicate that SWELL1 and PSA interact with each other in native cells. Treatment with hypotonic solution did not alter the IP levels of PSA, suggesting that their association is not dynamically regulated by cell swelling during VRAC activation (**Figure 1D**). To determine whether the SWELL1–PSA interaction is mediated indirectly through other VRAC subunits, we co-expressed both proteins in *LRRC8⁻/⁻* HEK293 cells, in which all five *LRRC8* genes are disrupted^14^. Reciprocal co-IP revealed a similar interaction between SWELL1 and PSA (**Figure 1E and F**), indicating that their association is independent of other LRRC8 subunits. Notably, the main mRNA transcript of PSA contains two AUG start codons^20^. Although the first AUG is typically assumed to serve as the translation initiation site, previous studies based on protein sequencing and antibodies have shown that PSA translation instead initiates from the second start codon^21,22^. Consistent with these findings, we observed that the longer PSA isoform containing an additional 44 N-terminal amino acids—translated from the first AUG— predominantly mislocalizes to mitochondria (**Figure S1**). Accordingly, all subsequent experiments employed a shorter PSA cDNA construct initiating at the second AUG. To examine the subcellular localization of SWELL1 and PSA, we generated HeLa cells with GFP inserted into the endogenous *SWELL1* locus using CRISPR-Cas9 and transfected them with a plasmid encoding a PSA-mCherry fusion protein. Live-cell imaging revealed that native SWELL1 is primarily localized to the plasma membrane as expected, while PSA is distributed throughout the cytosol (**Figure 1G**). Consistent with their interaction, partial co-localization of PSA and SWELL1 was observed near the plasma membrane (**Figure 1G and H**).

### Cryo-EM structure of the SWELL1–PSA complex

To gain structural insight into their interaction, we first purified human PSA and confirmed its binding to detergent-solubilized human SWELL1 by fluorescence-detection size exclusion chromatography (FSEC)^23^ (**Figure S2A**). We then purified SWELL1 and mixed with PSA for single-particle cryo-EM analysis (**Figure S2B and S2C**). The three-dimensional (3D) structure of the SWELL1–PSA complex was reconstructed to an overall resolution of 3.19 Å with C3 symmetry imposed (**Figure 2A** and **Figure S3**). To improve the local resolution of the PSA-bound SWELL1 LRR domains, symmetry expansion and focused refinement were applied, yielding a 3.74 Å map (**Figure S3-S5**; **Table S1**). The quality of the map enabled building of a structural model, with the exception of the N-terminus and two small, disordered regions within the first extracellular and intracellular loops of SWELL1 (**Figure S5**).

**Figure 2.**
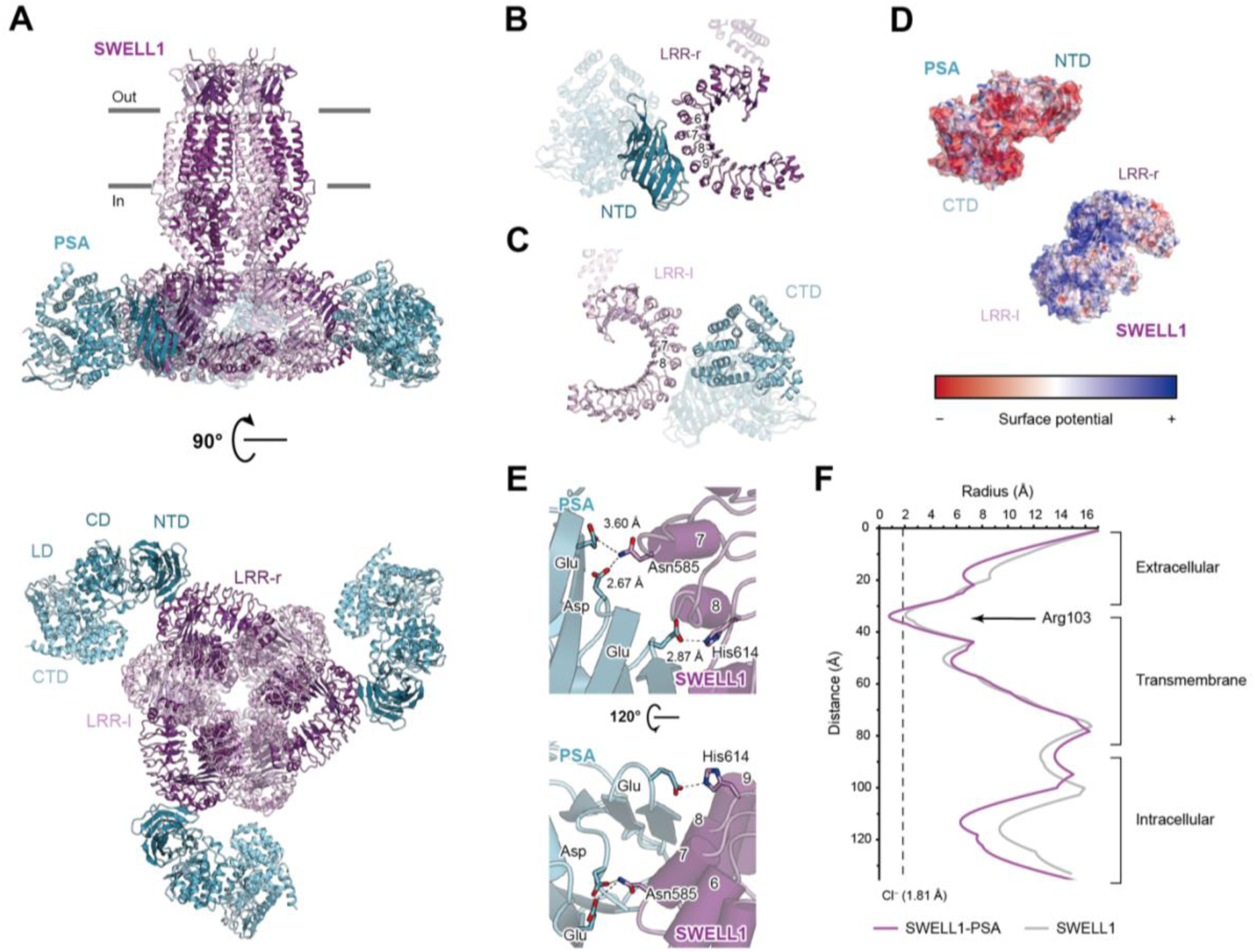
Cryo-EM structure of the SWELL1-PSA complex. (A) Ribbon model of the SWELL1–PSA complex, shown in side (top) and bottom (bottom) views. SWELL1 and PSA are colored magenta and blue, respectively, with the two SWELL1 subunits in each dimer distinguished by light and dark. Gray bars indicate the approximate boundaries of the lipid membrane. NTD: N-terminal domain; CD: catalytic domain; LD: linker domain; CTD: C-terminal domain (B and C) Binding site of PSA and the LRR domains, viewed from NTD side (B) and CTD side (C). The number of repeats contacted by PSA are labeled. (D) Surface potentials of PSA and the LRR domains. (E) Close-up view of the interface between NTD of PSA and LRR-r, viewed from the side (top) and from the top (bottom). LRRs contacted by PSA and key interacting amino acids are labeled. (F) Pore radii of PSA-bound SWELL1 (purple) and apo SWELL1 channels (gray, PDB ID: 5ZSU). The radius of dehydrated chloride ion is indicated as the black dashed line.

PSA adopts a V-shaped architecture in the complex structure, with minimal conformational changes relative to the previously reported apo form^24^ (Protein Data Bank (PDB) ID: 8SW0) (**Figure 2A**). It simultaneously engages the convex side of two adjacent LRR domains of SWELL1, interacting tightly with the right LRR (LRR-r) at repeats 6–9 via the N-terminal domain (NTD), burying a surface area of ∼700 Å^2^ (**Figure 2B**), and more loosely with the left LRR (LRR-l) at repeats 7–8 via the C-terminal domain (CTD), burying ∼193 Å^2^ (**Figure 2C**). 3D classification of the locally refined density maps revealed exclusively dimer classes of the LRR domains (**Figure S3**), suggesting that PSA binding stabilizes the LRR domains into a dimeric configuration and suppresses their flexibility^25–28^. The interacting surfaces of PSA and the LRR domains exhibit complementary electrostatic features, with PSA presenting predominantly negatively charged surfaces and the LRR domains displaying positively charged regions at the binding interfaces (**Figure 2D**). Consistently, at the interface between the NTD of PSA and LRR-r, three acidic side chains—D55, E81, and E107—are positioned opposite two polar residues from LRR-r (N585 and H614), likely forming hydrogen bonds (**Figure 2E**).

In the complex structure, the overall architecture of the homo-hexameric SWELL1 also closely resembles the previously reported C3-symmetric apo structures^25–29^, which are thought to represent a closed channel conformation. The root-mean-square deviation (RMSD) is less than 3 Å compared to one of the SWELL1 apo structures (PDB ID: 5ZSU) determined under the same 150 mM NaCl condition^27^ (**Figure S6A**). Notably, PSA binding slightly narrows the ion conduction pore at its most constricted site, formed by arginine 103 (R103), reducing the pore radius to less than that of a chloride ion (**Figure 2F** and **S6b**). Additionally, the helical regions of the intracellular loops lining the pore also shift modestly inward (**Figure 2F** and **S6a**), which may influence the propagation of conformational changes from the LRR domains to the pore region^30^. Nevertheless, the significance of these conformational changes in channel activation remains unclear considering previous structural studies of SWELL1 in complex with inhibitory and potentiating sybodies^2,18^.

### PSA broadly inhibits VRAC activation triggered by diverse stimuli

The flexibility of the LRR domains have been shown to couple to VRAC activation^2,3^. Given its unique binding mode, we next investigated whether PSA modulates VRAC gating. To this end, we transfected PSA into HEK293 cells and performed whole-cell patch-clamp recordings to measure endogenous VRAC currents activated by either extracellular hypotonicity-induced cell swelling or intracellular low ionic strength. Remarkably, PSA overexpression completely suppressed VRAC currents in response to both stimuli (**Figure 3A-E**). To ensure that this inhibition was not due to impaired channel trafficking, we performed surface biotinylation assays and observed comparable levels of SWELL1 at the plasma membrane in PSA-overexpressing and control cells (**Figure 3A**). Beyond cell swelling and low ionic strength, VRACs can also be activated—albeit to a lesser extent—by other more physiologically relevant stimuli, such as sphingosine-1-phosphate (S1P)^7^, an important inflammatory mediator. Although the activation mechanisms likely differ, PSA overexpression similarly suppressed S1P-induced VRAC currents in mouse BV2 microglial cells (**Figure 3F and 3G**). Together, these findings indicate that PSA broadly inhibits VRAC activation downstream of diverse stimuli.

**Figure 3.**
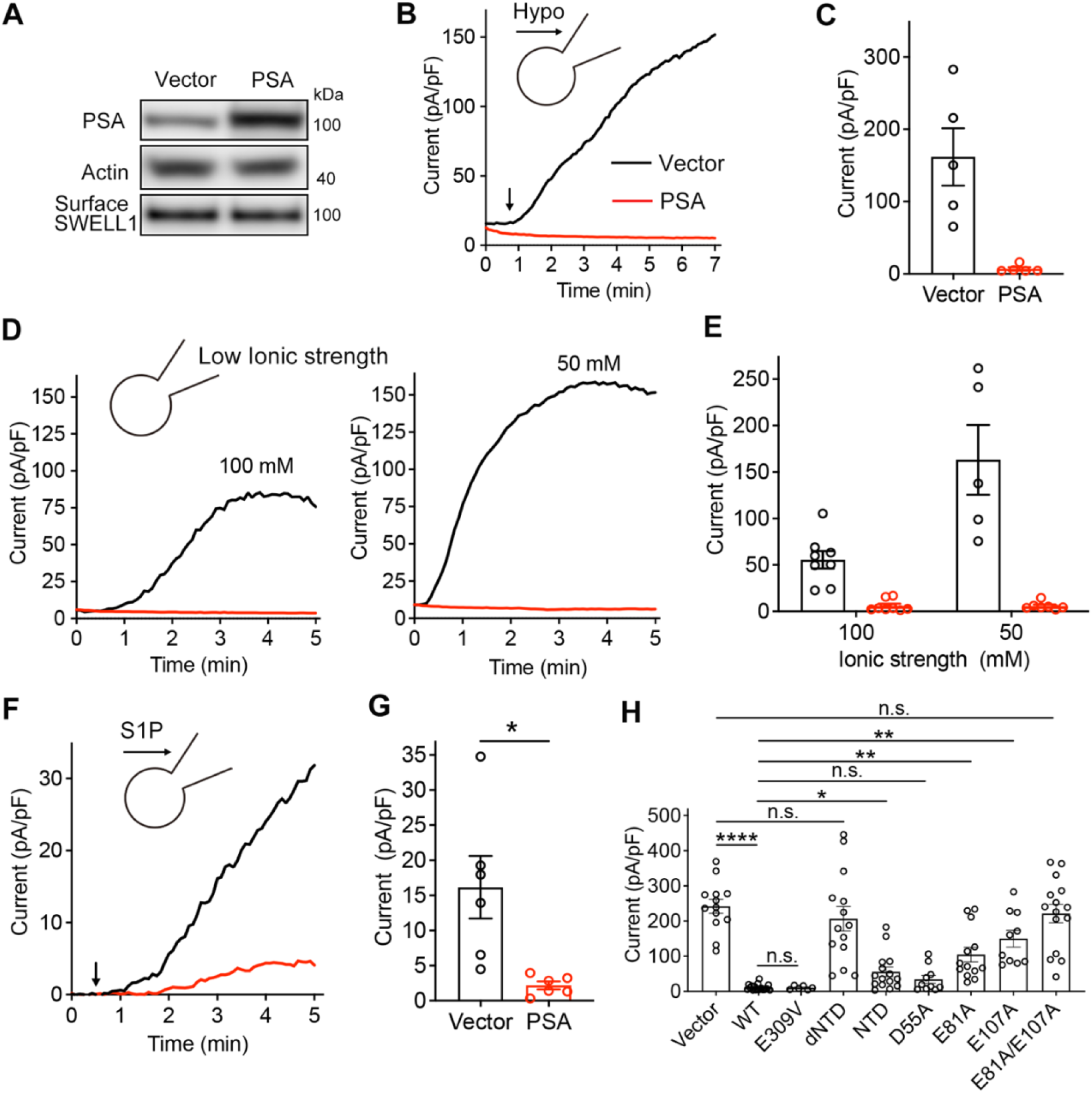
PSA broadly inhibits VRAC activation triggered by diverse stimuli. (A) Immunoblotting confirming PSA overexpression in HEK293 cells, showing no change in surface SWELL1 protein levels. (B and C) Time course (B) and quantification (C) of hypotonicity (230 mOsm/kg, applied at the arrow)-activated VRAC currents at +100 mV in HEK293 cells transfected with vector or PSA. (D and E) Time course (D) and quantification (E) of low intracellular ionic strength (100 mM, left; 50 mM, right)-activated VRAC currents at +100 mV in HEK293 cells transfected with vector or PSA. (F and G) Time course (F) and quantification (G) of background-subtracted 100 nM S1P-activated VRAC currents at +100 mV in BV2 microglial cells transfected with vector or PSA. Two-tailed *t*-test, *p*\* < 0.05. (H) Low intracellular ionic strength (50 mM)-activated VRAC currents at +100 mV in HeLa cells transfected with vector, WT PSA or various PSA mutants. One-way ANOVA and post hoc test, *p*\* < 0.05, *p* ** < 0.01, *p* **** < 0.0001, n.s., not significant. All bars represent mean ± SEM for the number of cells indicated.

To determine whether the aminopeptidase activity of PSA is required for VRAC inhibition, we treated HeLa cells with tosedostat, a potent inhibitor of M1 family aminopeptidases (**Figure S7A**). Tosedostat treatment had no effect on the ability of PSA overexpression to suppress VRAC currents (**Figure S7B**), indicating that PSA inhibits VRACs independently of its enzymatic activity. To further test this, we expressed a catalytically inactive PSA mutant in which the active-site glutamate was replaced with valine^31^ (E309V; also denoted as E353V in some publications where the longer PSA isoform was used) (**Figure S7C and S7D**). Like the wild-type (WT) protein, the E309V mutant robustly suppressed VRAC currents activated by low intracellular ionic strength (**Figure 3H**), further supporting that PSA’s enzymatic activity is dispensable for its inhibitory effect on VRACs. In contrast, deletion of the PSA NTD—the primary region interacting with SWELL1—completely abolished its ability to inhibit VRAC activity (**Figure 3H**), highlighting the essential role of the NTD in channel regulation. Consistent with this, the NTD alone was sufficient to suppress VRAC activity (**Figure 3H**), although it was less potent than the WT protein. This suggests that other domains, such as the CTD, also contribute to stabilizing the SWELL1– PSA interaction and enhancing channel inhibition.

To further dissect the SWELL1–PSA interface, we individually substituted three negatively charged residues in the NTD (D55, E81, and E107) with alanine. While the D55A mutant retained its inhibitory effect, both E81A and E107A were less potent in suppressing VRAC currents (**Figure 3H**). Strikingly, despite being expressed at similar levels as WT (**Figure S7D**), the E81A/E107A double mutant completely lost its ability to inhibit VRAC activity (**Figure 3H**). Taken together with structural data, these functional results support a model in which PSA inhibits VRAC gating by interacting with and stabilizing of the SWELL1 LRR domains.

### Deletion of PSA enhances basal VRAC activity

Given its ubiquitous expression, endogenous PSA may also play a regulatory role in VRAC activation. To test this hypothesis, we generated PSA knockout (KO) HeLa cells using CRISPR-Cas9 and performed whole-cell patch-clamp recordings (**Figure 4A**). Interestingly, PSA KO cells exhibited substantial basal currents under isotonic conditions (**Figure 4B and 4C**). These basal currents displayed hallmark features of VRACs, including outward rectification and sensitivity to dicumarol (**Figure 4B, S8A and S8B**), a known VRAC inhibitor^7^. Additional deletion of SWELL1 in PSA KO cells completely abolished the basal currents (**Figure 4B and 4C**), confirming that they are indeed mediated by VRACs. Similar basal VRAC currents were also observed in PSA KO human Jurkat T cells (**Figure 4E and 4F**), supporting a physiological role for PSA in maintaining VRACs in a closed state under resting conditions. Notably, hypotonicity-induced cell swelling further activated VRAC currents in PSA KO cells to levels comparable to those in control cells (**Figure 4D and 4F)**, indicating that PSA deletion does not enhance maximal VRAC activity. Moreover, knocking out PSA did not alter VRACs’ anion selectivity for iodide or glutamate as determined by reversal potential measurements (**Figure S8C and S8D**). These results are consistent with the model that cytosolic PSA functions as an auxiliary, but not a pore-forming, subunit of VRACs.

**Figure 4.**
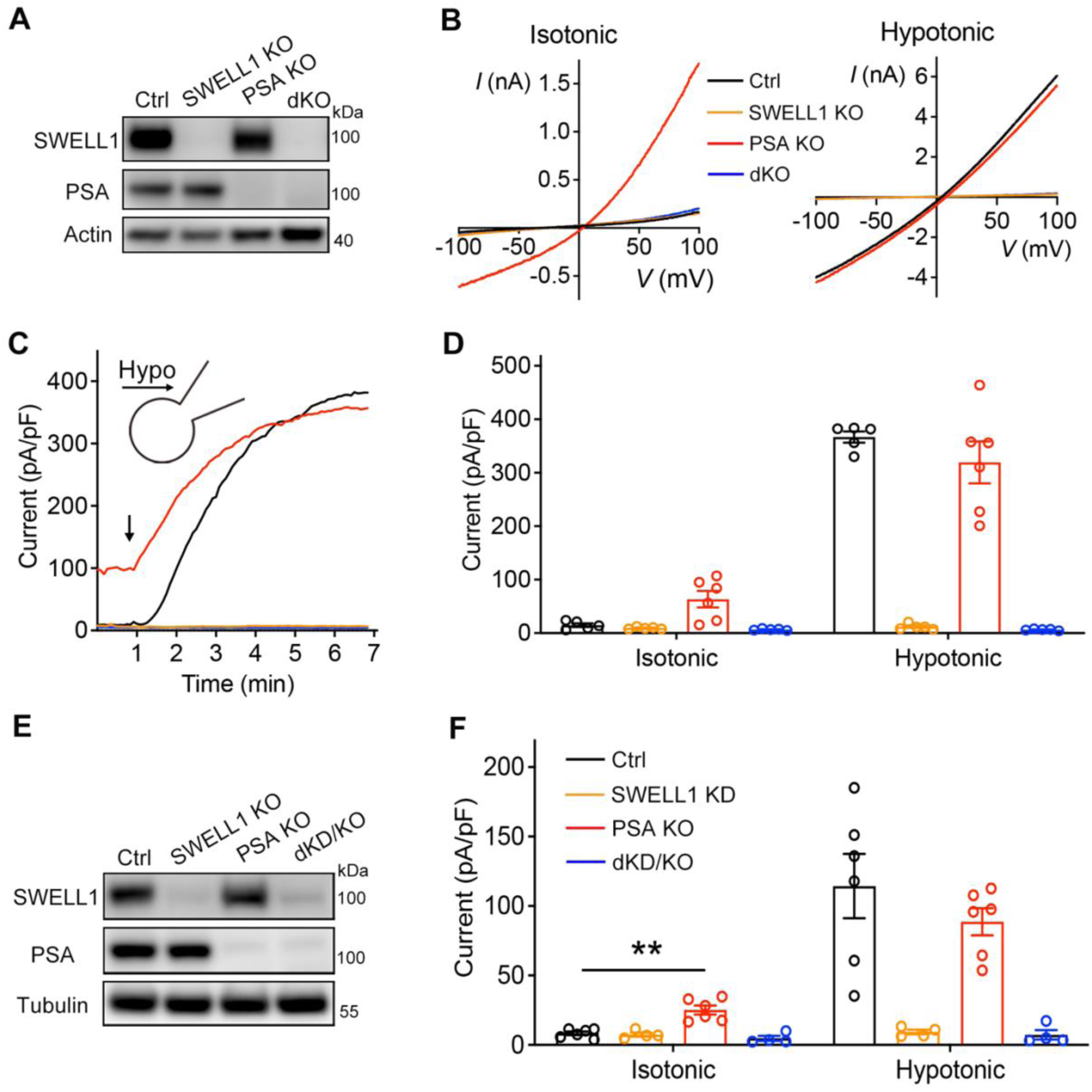
Deletion of PSA enhances basal VRAC activity. (A) Immunoblotting of protein expression in control, SWELL1 KO, PSA KO, and SWELL1/PSA double KO (dKO) HeLa cells. (B) Representative whole-cell currents recorded using a voltage ramp protocol for various HeLa cell lines in isotonic solution (300 mOsm/kg, left) and following perfusion with hypotonic solution (230 mOsm/kg, right). (C and D) Time course (C) and quantification (D) of whole-cell currents at +100 mV in various HeLa cell lines in isotonic solution and following perfusion with hypotonic solution. (E) Immunoblotting of protein expression in control, SWELL1 knockdown (KD), PSA KO, and SWELL1 KD/PSA KO (dKD/KO) Jurkat cells. (F) Quantification of whole-cell currents at +100 mV in various Jurkat cell lines in isotonic solution and following perfusion with hypotonic solution. One-way ANOVA and post hoc test, *p* ** < 0.01. All bars represent mean ± SEM for the number of cells indicated.

To quantitatively assess the effect of PSA on VRAC gating, we systematically varied intracellular ionic strength under isotonic conditions and recorded VRAC currents^32^. As expected, PSA KO HeLa cells exhibited robust basal currents at the physiological ionic strength of 150 mM, which progressively declined at 175 mM and became undetectable at 200 mM (**Figure 5A and 5B**). In contrast, lowering the ionic strength to 125 mM failed to activate VRACs in control WT cells but elicited additional currents in PSA KO cells (**Figure 5A and 5B**). Further reduction to 100 mM initiated VRAC activation in control cells and induced even larger currents in PSA- deficient cells (**Figure 5A and 5B**). At ionic strengths below 75 mM, both control and KO cells displayed maximal VRAC activation; however, the kinetics of current development were markedly faster in the absence of PSA (**Figure 5A-C**). PSA deletion significantly increased the sensitivity of VRACs to ionic strength, reducing the half-maximal effective concentration (EC₅₀) from 113 mM in control WT cells to 89 mM in PSA KO cells (**Figure 5B**). Together, these data suggest that endogenous PSA binding attenuates VRAC sensitivity to intracellular ionic strength, thereby fine-tuning channel activity and preventing excessive activation under physiological conditions.

**Figure 5.**
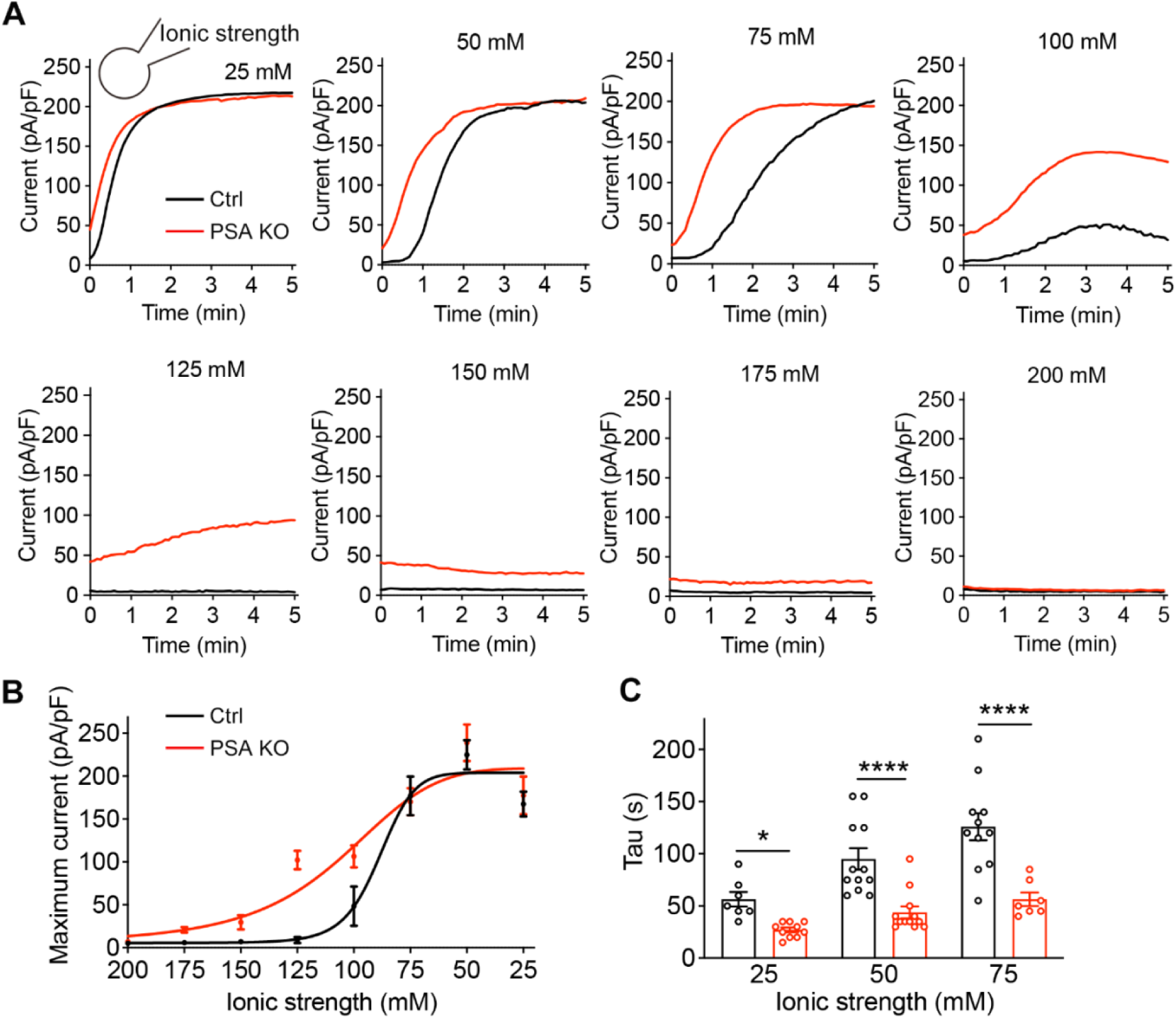
PSA reduces VRAC sensitivity to intracellular ionic strength. (A) Time course of low intracellular ionic strength-activated VRAC currents at +100 mV in control and PSA KO HeLa cells. (B) VRAC current-to-intracellular ionic strength relationship in control and PSA KO HeLa cells. (C) Time constant (tau) of the rising of VRAC currents activated by various low intracellular ionic strength in control and PSA KO HeLa cells. Two-way ANOVA with post hoc test, *p* * < 0.05, *p* **** < 0.0001. All bars represent mean ± SEM for the number of cells indicated.

### PSA regulates cGAMP transport by modulating VRAC activity

In addition to chloride ions, VRACs also permeate various small signaling molecules, including the immunomodulator cGAMP^4,10^, which is synthesized by the enzyme cGAS in response to cytosolic DNA. Extracellular cGAMP, released from damaged or diseased cells, can enter neighboring host cells through VRACs under physiological conditions, where it activates the innate immune STING pathway^4,33^. Given PSA’s role in modulating VRAC activity, we next examined whether PSA regulates cGAMP transport and downstream signaling. To this end, we treated Jurkat cells with extracellular cGAMP and measured STING phosphorylation (p-STING), the earliest detectable event following cGAMP import and binding. As expected, knockdown of SWELL1 reduced p-STING levels by ∼65% relative to total STING, consistent with VRAC’s role in cGAMP import (**Figure 6A and 6B**). In contrast, knocking out PSA markedly enhanced p-STING levels (**Figure 6A and 6B**), aligning with the elevated basal VRAC activity observed in these KO cells (**Figure 4E and 4F**). Notably, this enhancement was completely dependent on SWELL1 (**Figure 6A and 6B**), confirming that it occurs through the VRAC pathway. Similar results were also observed in TIME cells (**Figure 6C and 6D**), a telomerase-immortalized human microvascular endothelial (HMVEC) line, and HeLa cells (**Figure S9A and S9B**), indicating that PSA negatively regulates cGAMP transport through VRAC inhibition across diverse cell types. In contrast, PSA overexpression suppressed cGAMP-induced STING activation in HeLa cells (**Figure 6E and 6F**), further supporting its inhibitory effect on cGAMP transport. Consistent with the dispensable role of PSA’s aminopeptidase activity in VRAC regulation, treatment with tosedostat neither enhanced cGAMP uptake in control cells nor reversed the suppression of cGAMP transport by PSA overexpression (**Figure 6E and 6F**). Together, these results uncover a previously unrecognized role for PSA in regulating cGAMP transport and STING signaling through modulation of VRAC activity.

**Figure 6.**
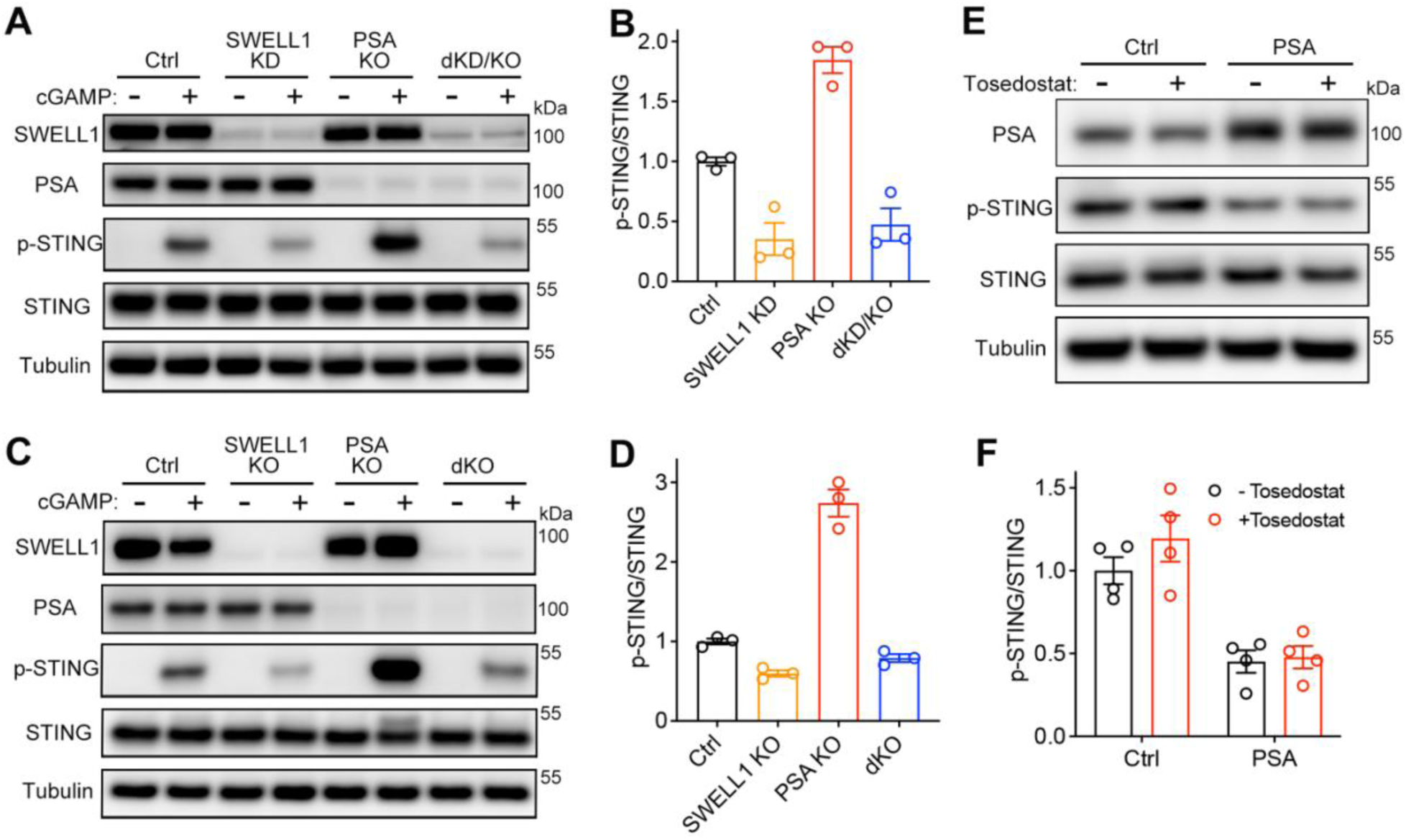
PSA regulates cGAMP uptake and STING signaling by modulating VRAC activity. (A and B) Immunoblot (A) and quantification (B) of phosphorylated STING (p-STING) and total STING in control, SWELL1 KD, PSA KO, and SWELL1 KD/PSA KO (dKD/KO) Jurkat cells treated with cGAMP (10 μM, 1 hr). p-STING/STING ratios were normalized to 1 in control cells. (C and D) Immunoblot (C) and quantification (D) of p-STING and total STING in control, SWELL1 KO, PSA KO, and SWELL1 KO/PSA KO (dKO) TIME cells treated with cGAMP (20 μM, 1 hr). p-STING/STING ratios were normalized to 1 in control cells. (E and F) Immunoblot (E) and quantification (F) of p-STING and total STING in control and PSA-overexpressing HeLa cells treated with cGAMP (50 μM, 1 hr), with or without tosdedostat (10 μM, 48 hrs including during cGAMP treatment). p-STING/STING ratios were normalized to 1 in control cells. All bars represent mean ± SEM for the number of experiments indicated.

## DISCUSSION

Through affinity purification and mass spectrometry, we have identified the ubiquitously expressed puromycin-sensitive aminopeptidase (PSA) as a novel auxiliary subunit that allosterically regulates VRAC activation. PSA binds to the convex surface of the SWELL1 LRR domains, reminiscent of the binding mode of three inhibitory sybodies^18^, particularly Sb1. However, unlike these sybodies, which interact with individual LRR domains, the V-shaped PSA simultaneously engages two adjacent domains—an elegant arrangement that likely further enhances its ability to stabilize the dimeric conformation of the LRR domains. Our finding that PSA inhibits VRAC activity further supports the notion that LRR domain mobility is coupled to VRAC activation^2,3^.

Although SWELL1 homo-hexameric channel exhibits poor activation properties and may not form under physiological conditions^25^, its structure is more homogenous than those of SWELL1/LRRC8 heteromeric channels, making it more amenable to structural interpretation of PSA binding. Importantly, the trimer-of-dimers architecture observed in a subset of SWELL1 homomers appears to be conserved in certain SWELL1/LRRC8C and SWELL1/LRRC8D heteromers, where the predominant configuration consists of two SWELL1 dimers and one LRRC8 dimer^34–36^. Consistent with the sequence and structural conservation among LRRC8 paralogues, we found that PSA also interacts individually with other LRRC8 subunits when co-expressed in *LRRC8⁻/⁻* cells (**Figure S10**). While its binding affinities to different LRRC8 subunits may vary and remain to be determined, the binding mode of PSA observed in the SWELL1 homomers is likely conserved in the heteromeric channels. In addition to allosterically modulating VRAC activity, PSA may also facilitate channel assembly by stabilizing LRRC8 dimers as discrete structural units during biogenesis. These hypotheses will be tested in future structural and functional studies, for which the current work provides an important foundation.

Our results suggest a model in which some pore-forming subunits of VRACs (SWELL1 and/or other LRRC8 proteins) are bound and stabilized by endogenous PSA in native cells (**Figure 7**). This interaction reduces the sensitivity of the channels to ionic strength and likely other stimuli, thereby preventing excessive basal VRAC activity under normal physiological conditions. When PSA is overexpressed or upregulated, it saturates all available binding sites, resulting in maximal stabilization of the LRR domains and rendering the channels unresponsive to various stimuli. Our findings raise an intriguing possibility that, during channel activation, the inhibitory PSA auxiliary subunit dissociates from VRACs as part of the gating mechanism. However, this scenario appears to be unlikely, as the SWELL1–PSA interaction is not disrupted by cell swelling and the associated decrease in intracellular ionic strength (**Figure 1D**). Furthermore, VRAC currents are readily reversible upon removal of stimuli, suggesting a dynamic gating mechanism that does not require complete dissociation of PSA. Finally, the interaction between PSA and the LRR domains is primarily electrostatic and is likely strengthened, rather than weakened, under the low ionic strength conditions induced by cell swelling. Nevertheless, it remains possible that subtle changes in PSA binding mode may increase LRR domain flexibility, thereby contributing to VRAC activation.

**Figure 7.**
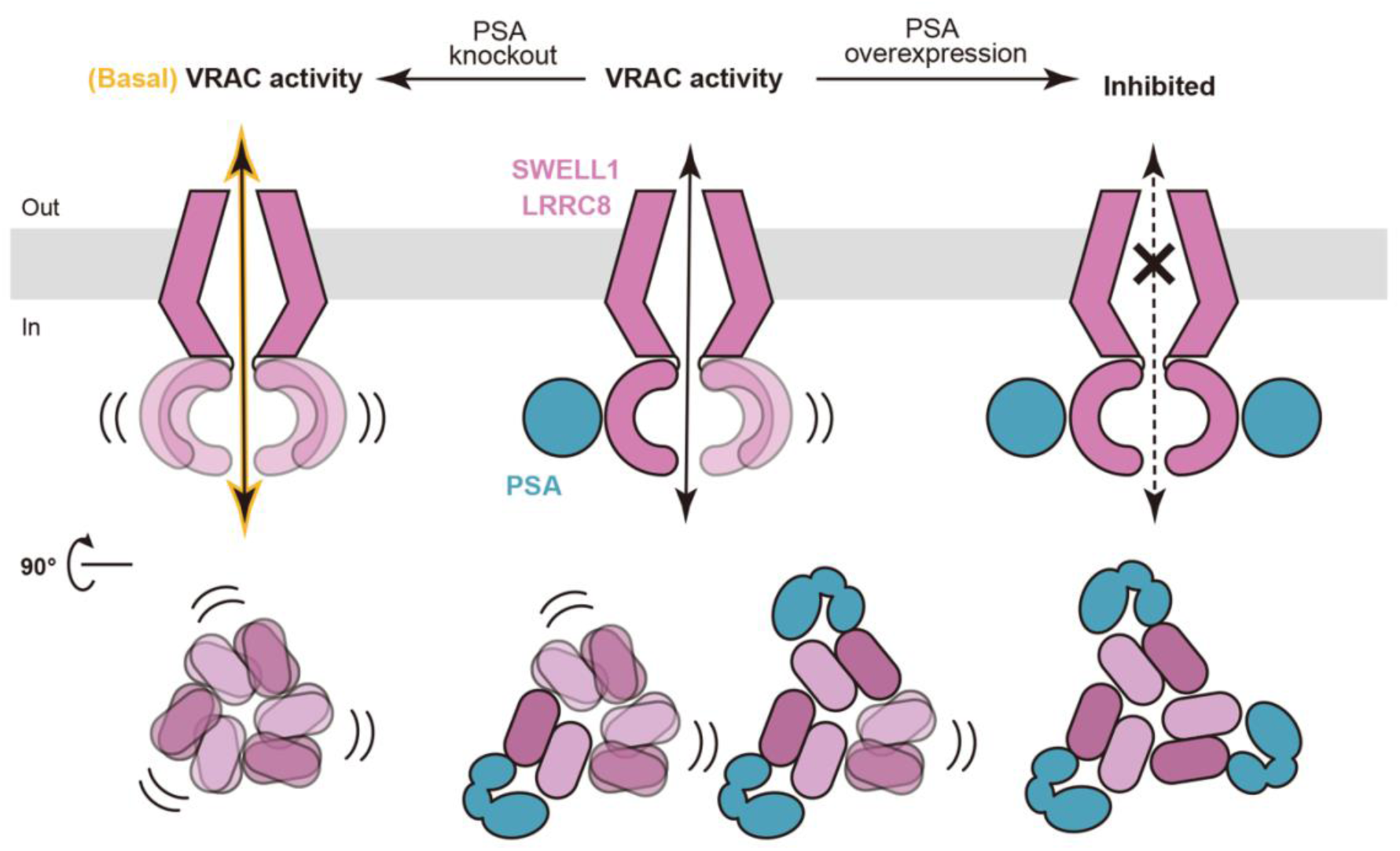
Model for allosteric modulation of VRACs by PSA. Schematic illustrating how PSA binding to the intracellular LRR domains of SWELL1/LRRC8 stabilizes a dimeric configuration, restricting their mobility and thereby inhibiting VRAC activation. This allosteric regulation reduces the channel’s sensitivity to various stimuli, keeping VRACs closed or less active under resting conditions (middle). Deletion of PSA increases LRR domain flexible and elevates basal VRAC activity (left), whereas PSA overexpression fully stabilizes the LRR domains and suppresses channel activation (right).

Our results indicate that PSA acts as a ubiquitous regulator of VRAC activity, in contrast to typical auxiliary subunits, which are expressed in a tissue-specific manner to modulate channel function in select cell types. Accordingly, beyond its canonical function in proteolysis, PSA may also participate in the diverse physiological and pathological processes regulated by VRACs. Indeed, we demonstrate that PSA modulates cGAMP import across multiple cell types. Thus, it may represent a novel therapeutic target for enhancing or restricting extracellular cGAMP uptake and STING signaling, with potential to boost anti-tumor immunity or mitigate pathological inflammation in a range of diseases^37^. Interestingly, a recent study identified PSA as a driver of cisplatin resistance in human bladder cancer cells by regulating intracellular cisplatin levels^38^. Depletion of PSA sensitized resistant cells to cisplatin, whereas its overexpression in sensitive cells conferred increased resistance. In follow-up work, the authors showed that PSA interacts with VRACs and regulates cisplatin import in a SWELL1-dependent manner^19^. Through pharmacological and mutagenesis approaches, they concluded that PSA’s enzymatic activity is essential for mediating cisplatin resistance and proposed its pharmacological inhibition as a potential strategy to improve patient responses to cisplatin-based chemotherapy^19,38^. Our work complements their findings by providing the structural basis of the SWELL1–PSA interaction and elucidating the molecular mechanism by which PSA regulates VRAC activity. However, our study indicates that PSA’s enzymatic activity is dispensable for this regulatory function. The reason for this discrepancy remains unclear. One possibility is that tosedostat may target proteins other than PSA in bladder cancer cells, contributing to the observed cisplatin sensitization. Additionally, the catalytically inactive mutant used in their study was generated in the longer PSA isoform, which predominantly localizes to mitochondria and therefore may behave differently from the corresponding mutant expressed from the shorter isoform used in our experiments. Clarifying this discrepancy will be important, as our findings suggest that disrupting the physical interaction between SWELL1 and PSA, rather than inhibiting PSA’s enzymatic activity, may represent a more effective therapeutic strategy to enhance VRAC activity and, in turn, increase sensitivity to cisplatin-based chemotherapy or cGAMP-mediated anti-tumor immunity.

## METHODS

### Mass spectrometry

HEK293 cells were transfected with either pCMV vector or pCMV-SWELL1-Flag (OriGene, RC208632) using Lipofectamine 2000 transfection agent (Life Technologies) for 2 days prior to lysis in radioimmunoprecipitation assay (RIPA) buffer (50 mM Tris-HCl, pH 7.4, 150 mM NaCl, 2.5% deoxycholic acid, 10% NP-40, 10 mM EDTA) containing 1% protease inhibitor cocktail (Roche). Lysates were rotated at 4°C for 30 min and clarified by centrifugation at 13,800 × g for 30 min. The resulting supernatants were incubated with anti-Flag M2 magnetic beads (Sigma) overnight at 4°C. The beads were washed four times for 5 min each and eluted with 50 µg/ml 3× Flag peptide (Sigma) in a buffer containing 20 mM Tris, pH7.5, 150 mM NaCl, 0.05% DDM and a cocktail of protease inhibitors. The affinity-purified samples were separated by SDS–PAGE gel electrophoresis and visualized by Coomassie blue staining. To increase proteomics coverage, the entire gel lane from samples purified from control and SWELL1-Flag-expressing cells was excised into 20 evenly sized slices, each subjected individually to mass spectrometry before pooling for data analysis. Gel bands were de-stained, reduced with dithiothreitol (DTT), alkylated with iodoacetamide, digested with trypsin overnight, and dried under vacuum. Dried samples were reconstituted in 0.1% formic acid and analyzed on a Q-Exactive Orbitrap Mass Spectrometer as previously reported^16^. Data files were imported and automatically aligned using Mascot. Peptide false discovery rate (FDR) was set to 1%, and peptide identification was performed against a human slice of the UniProt database. Proteins with more than 3 unique peptides detected in affinity-purified SWELL1-Flag samples but absent from controls were selected as positive hits.

### Molecular Biology

The coding sequence for the longer human PSA isoform (NM_006310) with Myc and Flag tags in pCMV vector was obtained from OriGene (RC209037). The coding sequence starting from the second ATG (2,625 bp) with or without the tags was PCR amplified and sub-cloned into pIRES2-EGFP (Clontech) and pLenti-EF1a (Addgene). The PSA point mutations and truncations were introduced using Site-Directed Mutagenesis Kit (NEB). Myc-Flag-tagged cDNAs of human SWELL1 (NM_019594), LRRC8B (NM_015350), LRRC8C (NM_032270), LRRC8E (NM_025061) in pCMV vector were obtained from OriGene (RC208632, RC205553, RC222603, RC209849). The coding sequences without a tag were PCR amplified and sub-cloned into pIRES2-EGFP vector (Clontech). The cDNA of human LRRC8D (NM_018130) in pCMV6-Entry vector was obtained from Amsbio(SC319580).

### Cell culture

HeLa cells, HEK293 cells, and BV2 microglial cells were maintained in Dulbecco’s Modified Eagle’s Medium (DMEM) supplemented with 10% fetal bovine serum (FBS) and 1% penicillin/streptomycin. SWELL1-Flag knock-in HeLa cells^16^, SWELL1 KO HeLa cells^5^, and SWELL1 KO BV2 cells^7^ were generated previously. *LRRC8⁻/⁻* HEK293 cells were generously provided by T. Jentsch^14^. Jurkat T cells were grown in Roswell Park Memorial Institute (RPMI) 1640 medium containing GlutaMAX, supplemented with 10% FBS. SWELL1 knockdown Jurkat cells were generated using lentiviruses expressing SWELL1-specific shRNA (GGUACAACCACAUCGCCUA) as previously reported^15^. Control and SWELL1 KO telomerase-immortalized human microvascular endothelial (TIME) cells (generously provided by Dr. Linying Li) were cultured in vascular cell basal media supplemented with a microvascular endothelial cell growth kit-VEGF (ATCC) and 1% penicillin/streptomycin.

### Immunoblot and antibodies

Homogenates of cultured cells were prepared in RIPA buffer containing 1% protease inhibitor cocktail (Roche). Lysates were rotated at 4°C for 30 min and clarified by centrifugation at 13,800 × g for 30 min. The total protein levels were determined by BCA protein assay kit (Thermo scientific). 6 × Laemmli SDS-sample buffer (Boston bioproduct) was added into the supernatants then followed by boiling at 95°C for 10 min. Samples were resolved on 4–20% Bis-Tris SDS-PAGE gels (Life Technologies) and transferred to Immobilon-P PVDF membranes (Millipore). Membranes were blocked in Tris-buffered saline (TBS) containing 0.05% Tween-20 (TBST) and 5% fat-free milk for 1 h at room temperature (RT) before incubation with primary antibodies. Membranes were probed overnight at 4°C with the following primary antibodies: anti-PSA (Invitrogen, #MA5-47084), anti-SWELL1^5^, anti-phospho-STING (Ser366) (Cell Signaling Technology, #19781), anti-STING (Cell Signaling Technology, #13647), anti-Myc (Cell Signaling Technology, # 5605), anti-alpha-Tubulin (Cell Signaling Technology, #3873) and anti-beta-Actin (Cell Signaling Technology, #4967). After primary antibody incubation, membranes were washed by TBST for three times at RT and incubated with horseradish peroxidase (HRP)-conjugated secondary antibodies: Goat Anti-Rabbit IgG (H+L) (Jackson ImmunoResearch Laboratories, #111-035-144) or Goat Anti-Mouse IgG (H+L) (Jackson ImmunoResearch Laboratories, #111-035-166) diluted in 5% fat-free milk for 1 h at RT. The membranes were washed by TBST for three times and 10 min each at RT. Immunoreactive bands were visualized using enhanced chemiluminescence (ECL). Films were scanned with the AI-680 system (GE) and analyzed using ImageJ.

### Immunoprecipitation

For IP of endogenous proteins, HeLa SWELL1-Flag knock-in cells were crosslinked with 0.5% formaldehyde in washing buffer (20 mM HEPES, pH 7.6, 10 mM KCl, 1.5 mM MgCl_2_) for 10 min on ice^39^. The reaction was quenched by adding one-fourth volume of 1.25 M glycine for 10 min. The cells were centrifuged at 1,000 × g for 5 min at 4°C and resuspended with washing buffer. After sonication with 10-sec pulses for 2 min on ice, one-ninth volume of 5 M NaCl was added to the extract. The lysates were centrifuged at 13,800 × g for 10 min at 4°C, and the resulting supernatants were incubated overnight at 4°C with anti-Flag monoclonal antibody (Sigma, F1804) or anti-PSA polyclonal antibody (Invitrogen, #PA5-83788) with gentle mixing. After incubated with antibodies, the protein G magnetic beads were pre-washed with TBS buffer and were added into the supernatant-antibody mixes followed with gently rotating for 2 h at 4°C. Precipitated proteins were washed using TBS buffer for four times. To elute the proteins from beads, 2× Laemmli SDS-sample buffer (Boston bioproduct) was added followed by boiling at 95°C for 10 min. The affinity-purified samples were analyzed via immunoblot as described above. IgG isotype was used a negative control, and whole-cell lysates served as input controls. For some experiments, cells were treated with hypotonic solution (230 mOsm kg^−1^, as described in the electrophysiological recordings) for 0, 5, or 30 min prior to IP.

For IP of overexpressed proteins, SWELL1 and PSA (only one of which has a Flag tag) were co-transfected in *LRRC8⁻/⁻* HEK293 cells using Lipofectamine 2000 transfection reagent (Life Technologies). After 2 days, cells were lysed in RIPA buffer containing 1% protease inhibitor cocktail. Lysates were clarified by centrifugation at 13,800 × g for 30 min. IP was performed using anti-Flag-M2 magnetic beads (Sigma) with overnight incubation at 4°C under gentle mixing. Precipitated proteins were washed, eluted, and analyzed via immunoblot as described above.

### Surface biotinylation

Surface biotinylation was performed as previously described^40^. One day after PSA transfection, HEK293 cells were rinsed once with ice-cold PBSCM (PBS containing 0.1 mM CaCl₂, 1 mM MgCl₂, pH 8.0), then incubated with Sulfo-NHS-SS-biotin (1 mg ml^−1^; Thermo Scientific) for 20 min at 4°C. Unreacted biotin was quenched by washing cells twice for 5 min with 20 mM glycine in PBSCM. Cells were lysed in buffer containing PBS, 50 mM NaF, 5 mM sodium pyrophosphate, 1% NP-40, 1% sodium deoxycholate, 0.02% SDS, and the protease inhibitor cocktail. Equal amounts of protein were incubated overnight at 4 °C with NeutrAvidin agarose beads (Thermo Scientific), followed by four washes with lysis buffer. Biotinylated proteins were eluted in 2× SDS loading buffer by boiling at 70 °C for 10 min, then analyzed by SDS-PAGE and immunoblot with anti-SWELL1 antibody and anti-Tubulin as a negative control.

### Generation of PSA knockout cells

To generate PSA knockout (KO) cells, we used CRISPR-Cas9 technology^41^. The guide RNA (5’-GGCCAAACTAAAAATTCTAA-3’, sense strand) targeting PSA was cloned into PX458-mCherry (modified from PX458-GFP from Addgene) and transfected in HeLa cells or SWELL1 KO HeLa cells using Lipofectamine 2000. 2 days post-transfection, single mCherry-positive cells were FACS-sorted into 96-well plates. After 2-3 weeks, single KO colonies were isolated based on immunoblot and target-site-specific PCR followed by Sanger-sequencing. For hard-to-transfect cells (i.e., Jurkat and TIME cells), the same guide RNA was cloned into lentiCRISPR-v2-Blast vector (Addgene). Lentiviruses were produced by co-transfecting it with packaging vectors (pVSV-G, pMDL, and pRSV) in HEK293 cells following Addgene’s instruction. Jurkat and TIME cells (both control and SWELL1 knockdown or knockout) were transduced in the presence of 8 μg/mL polybrene, selected with blasticidin (2.5-10 μg/mL) 3 days post-transduction for 2-3 weeks. The panel of KO cells was validated by immunoblot.

### Live cell imaging

SWELL1-GFP Knock-in HeLa cells was generated by CRISPR-Cas9-mediated homologous recombination. The guide RNA (5’-GTGCTGGGCCGGCCTCGCTC-3’, antisense strand) targeting the sequence encoding C-terminus of SWELL1 was cloned into PX458-mCherry vector and was co-transfected to HeLa cells with a pBluescript plasmid carrying GFP and two homology arms. 2 days post-transfection, single mCherry-positive cells were FACS-sorted into 96-well plates After 2-3 weeks, single clones of SWELL1-GFP knock-in HeLa cells were selected based on target-site-specific PCR and TA cloning followed by Sanger-sequencing. The knock-in cells were plated onto glass-bottom dishes and transfected with pLenti-EF1a-PSA-mCherry for one day before imaging. For the mitochondria co-localization study, HeLa cells were co-transfected with pLenti-EF1a-Long-PSA-mCherry and Su9-EGFP (Addgene). Live cells were imaged on a Zeiss LSM900 confocal microscope 24 hours post-transfection. Images were processed using custom macros in ImageJ FIJI.

### Protein expression and purification

The shorter human PSA isoform was PCR amplified from pCMV-PSA and inserted into the pFastBac vector (Gibco), with a thrombin cleavage site followed by mCherry and Strep-tag II at C-terminus. Baculoviruses were generated in *Spodoptera frugiperda* ExpiSf9 cells (Gibco), cultured in ExpiSf CD medium (Gibco) at 27°C. P2 baculovirus was used to infect ExpiSf9 cells at a density of approximately 3.5-5.0 × 10^6^ cells ml^−1^. After 84-96 h, the cells were collected by centrifugation (5,000 × *g*, 10 min, 4°C), frozen in liquid nitrogen, and stored at −80 °C until further purification. The cell pellet was disrupted by sonication in lysis buffer (20 mM Tris, pH 7.5, 150 mM NaCl, 10% glycerol). The cell debris was removed by centrifugation (2,000 × *g*, 10 min, 4°C) and the membrane fraction was removed by ultracentrifugation (142,000 × *g*, 1 h, 4°C). The supernatant was incubated with Strep-Tactin Superflow beads (IBA Lifesciences) for 1.5 h at 4°C. The protein-bound beads were poured into an open column and washed with 10 column volumes (CV) of lysis buffer. The proteins were eluted with elution buffer (100 mM Tris, pH 8.0, 150 mM NaCl, 2.5 mM desthiobiotin). The eluate was dialyzed in lysis buffer overnight with thrombin to digest the mCherry-Strep tag II. The dialyzed proteins were concentrated with a centrifugal filter unit (Merck Millipore, 50 kDa molecular weight cutoff) and purified by anion exchange chromatography on a HiTrap Q HP column (Cytiva), equilibrated with AE buffer (20 mM Tris, pH 7.5, 75 mM NaCl). The peak fractions were collected and dialyzed in dialysis-P buffer (20 mM Tris, pH 7.5, 150 mM NaCl) overnight. The dialyzed proteins were concentrated to 30 mg ml^−1^ with a centrifugal filter unit (Merck Millipore, 100 kDa molecular weight cutoff), ultracentrifuged at 98,000 × *g* for 25 min to remove the aggregation, frozen in liquid nitrogen, and stored at −80 °C until further use.

The cDNA of human SWELL1 was PCR amplified from pCMV-SWELL1 and inserted into the pFastBac vector, with a thrombin cleavage site followed by GFP and Strep-tag II at C-terminus. Baculoviruses were generated in *Spodoptera frugiperda* ExpiSf9 cells (Gibco), cultured in ExpiSf CD medium (Gibco) at 27°C. P2 baculovirus was used to infect ExpiSf9 cells at a density of approximately 3.5-5.0 × 10^6^ cells ml^−1^. After 84-96 h, the cells were collected by centrifugation (5,000 × *g*, 10 min, 4°C), frozen in liquid nitrogen, and stored at −80 °C until further purification. The cell pellet was resuspended and solubilized for 1 h and 4°C in solubilization buffer (20 mM Tris, pH 7.5, 150 mM NaCl, 10% glycerol, 1% n-dodecyl-β-D-maltoside (DDM; Anatrace), 0.2% cholesteryl hemisuccinate Tris salt (CHS; Anatrace)). Insoluble materials were removed by ultracentrifugation (186,000 × *g*, 1 h, 4°C). The detergent-soluble fraction was incubated with Strep-Tactin XT 4Flow beads (IBA Lifesciences) for 2.5 h at 4°C. The protein-bound beads were poured into an open column and washed with wash buffer (20 mM Tris, pH 7.5, 150 mM NaCl, 10% glycerol, 0.03% glyco-diosgenin (GDN; Anatrace)). The proteins were eluted with elution-B buffer (20 mM Tris, pH 7.5, 150 mM NaCl, 10% glycerol, 0.03% glyco-diosgenin (GDN; Anatrace), 50 mM biotin). The eluate was dialyzed in wash buffer overnight with thrombin to digest the mCherry-Strep tag II. The dialyzed proteins were concentrated with a centrifugal filter unit (Merck Millipore, 100 kDa molecular weight cutoff) and incubated with purified PSA in the ratio of 1:1 (w/w) for 1 h. The sample was then purified by size-exclusion chromatography (SEC) on a Superose 6 Increase 10/300 GL column (Cytiva), equilibrated with SEC buffer (20 mM Tris, pH 7.5, 150 mM NaCl, 0.03% GDN). The peak fractions were collected, concentrated to 3.5-4.0 mg ml^−1^ with a centrifugal filter unit (Merck Millipore, 100 kDa molecular weight cutoff), ultracentrifuged at 98,000 × *g* for 25 min to remove the aggregation, and immediately used for cryo-EM grid preparation.

### Cryo-EM image acquisition and data processing

C-flat (CF-1.2/1.3-2Au-50, Electron Microscopy Sciences) grids were coated with 50 nm Au by Sputter Coater Leica EM ACE600 and plasma cleaned with Ar/O_2_ by Tergeo Plasma Cleaner (Pie Scientific) to make holey 1.2/1.3 gold grids with gold mesh based on published methods^42,43^. A 3 µl portion of the protein sample was applied to a glow-discharged homemade gold grid, blotted using a Vitrobot Mark IV (FEI) under 4°C and 100% humidity conditions, and then frozen in liquid ethane. The grid images were obtained with a Glacios microscope (Thermo Fisher Scientific) operated at 200 kV and recorded by a Falcon 4i direct electron detector (Thermo Fisher Scientific). A total of 11,663 movies were obtained in the electron counting mode, with a physical pixel size of 1.1896 Å pixel^−1^. The data set was acquired with the EPU software, with a defocus range of −0.6 to −1.6 μm. Each image was dose-fractionated to 40 frames to accumulate a total dose of 40 e^−^ Å^−2^.

CryoSPARC (version 4.6.0) was preliminarily used for all aspects of data processing^44^. 3,117,293 particles were selected by blob-based auto-picking and extracted in 1.1896 Å pixel^−1^ from motion-corrected and dose-weighted micrographs. After iterative two-dimensional classification, 240,320 good particles were selected. Well-aligned 79,371 particles were selected from them and used to generate an initial 3D map. Initially selected 240,320 particles were subsequently refined using non-uniform refinement^44^ with C3 symmetry, resulting in an overall map at 3.19 Å. Focus masks for the channel core region (extracellular, transmembrane, and intracellular regions) and for LRR-PSA region (two LRRs and one PSA) are created separately and used for local refinement. Channel core region is refined with C3 symmetry, resulting in a local map at 3.16 Å. For the local refinement of LRR-PSA region, the particle stack was expanded by symmetry expansion with C3 symmetry. Expanded particles then undergo particle subtraction with the inverted mask of LRR-PSA region and 3D classification with the focus mask. 221,271 well-aligned particles are used for the local refinement, resulting in a local map at 3.74 Å (**Extended Data** Figs. 3**, 4**).

### Model building

The cryo-EM structure of homo-hexameric human SWELL1 (PDB: 7XZH) and the crystal structure of human PSA (PDB: 8SW0) were used as initial model and fitted into density map in COOT^45^. The model was refined using PHENIX with secondary structure restraints^46^ and modified manually in COOT iteratively. The electrostatic potential calculation was performed by the program APBS^47^. The van der Waals radii of the ion pathway were calculated using the HOLE program^48^. The figures depicting the molecular structures were prepared with UCSF ChimeraX^49^ and CueMol (http://www.cuemol.org/).

### Patch clamp electrophysiology

Patch clamp electrophysiology was performed as previously described^5,7^. Cells were seeded onto 12-mm poly-L-lysine–coated glass coverslips one day prior to whole-cell patch clamp recordings. In select experiments, cells were transfected with pIRES2-EGFP expressing WT PSA or various PSA mutants, for 24-36 h before recordings. For hypotonicity-activated VRAC current recordings, whole-cell patch clamp configuration was established in an isotonic bath solution containing (in mM): 105 NaCl, 2 KCl, 1 MgCl₂, 1 CaCl₂, 10 HEPES, and 80 mannitol (pH 7.3 adjusted with NaOH; osmolality ∼310 mOsm kg^−1^), and then a hypotonic solution that has the same ionic composition but without mannitol (osmolality ∼230 mOsm kg^−1^) was perfused. Recording electrodes (2–4 MΩ) were filled with an intracellular solution containing (in mM): 145 CsCl, 10 HEPES, 4 Mg-ATP, 0.5 Na₃-GTP, and 5 EGTA (pH 7.3 adjusted with CsOH; osmolality ∼300 mOsm kg^−1^). For S1P-activated VRAC current recordings in BV2 cells, everything was the same except that the cells were perfused with isotonic solution with 100 nM S1P.

For low intracellular ionic strength-activated VRAC current recordings, whole-cell patch clamp configuration was established in the same isotonic bath solution as above. Intracellular solutions contained (in mM) 25–200 CsCl in addition to 10 HEPES, 4 Mg-ATP, 0.5 Na₃-GTP, and 5 EGTA (pH 7.3 adjusted with CsOH). For intracellular solutions containing 25–125 mM CsCl, mannitol was added to adjust the osmolality to ∼300 mOsm kg^−1^; for intracellular solutions with 175 or 200 mM CsCl, mannitol was added to the bath solution to match its osmolality to that of the intracellular solution.

To determine VRAC permeability to Cl⁻, I⁻, and glutamate, HeLa cells were patched with Cl⁻-based intercellular solution containing (in mM): 120 NaCl, 10 HEPES, and 100 mannitol. Cells were bathed initially in an isotonic solution containing (in mM): 120 NaCl, 10 HEPES, and 100 mannitol, followed by perfusion with three different hypotonic solutions sequentially containing (in mM): 120 NaX, 10 HEPES, and 50 mannitol (where X = Cl⁻, I⁻, or glutamate). Solutions were adjusted to pH 7.3 with NaOH.

Recordings were performed using a MultiClamp 700B amplifier and a Digidata 1440A digitizer (Molecular Devices). Constant voltage ramps (every 5 s, 500 ms duration) were applied from a holding potential of 0 mV to ±100 mV. Data acquisition was carried out with Clampex 10.7 software (Molecular Devices).

### Aminopeptidase activity assay

HEK293 cells were lysed for 10 min on ice in digitonin-based lysis buffer containing (in mM): 20 HEPES (pH 7.6), 10 KCl, 1.5 MgCl₂, and 0.025% digitonin (Sigma). Lysates were then centrifuged at 1,000 × g for 10 min at 4°C. The supernatant was collected, and protein concentration was determined using the BCA Protein Assay Kit (Thermo scientific). To measure PSA aminopeptidase activity, 5 μg of total protein from each sample was incubated with the fluorescent substrate Leu-AMC (Bachem) at a final concentration of 100 μM in 96-well plates at 37°C for 1 h. Liberated AMC fluorescence was measured using an Infinite M Plex Microplate

Reader (TECAN) with excitation/emission wavelengths of 380/460 nm. The lysis buffer alone served as the blank control. The specificity of the assay was validated by using lysates from PSA KO HEK293 cells and through treatment with the PSA inhibitor tosedostat (10 μM).

### cGAMP treatment

Jurkat cells were seeded in 24-well plates at 2 × 10^6^ cells per well one day before treatment. Following removal of the old media by centrifugation (350 × g for 5 min), the cells were resuspended in the fresh media containing 10 μM cGAMP (Invivogen). HeLa cells and TIME cells were seeded in 6-well plates at 3 × 10^5^ cells per well one day before treatment. The old media was gently removed and replaced with fresh media containing cGAMP at a final concentration of 50 μM for HeLa cells and 20 μM for TIME cells. The cells were treated with cGAMP for 1 h prior to lysis and subsequent immunoblot analysis. The relative levels of phospho-STING versus total STING served as an indicator of cGAMP import. HeLa cells stably overexpressing PSA were generated by transducing lentiviruses packaged from pLenti-EF1a-PSA-P2A-GFP. 2 days post-transduction, GFP-positive cells were FACS-sorted. PSA overexpression in the stable cells were confirmed by immunoblot.

### Statistical analysis

All statistical analyses were performed using GraphPad Prism 9.1 software. Parametric tests, including paired or unpaired two-tailed *t*-tests, were used for comparisons between two groups with normally distributed data. For comparisons among more than two groups, analysis of variance (ANOVA) and post hoc tests was applied. Data are presented as means ± SEM. Statistical significance was defined as *P* < 0.05. The number of cells or independent experiments for each analysis is indicated in the corresponding figure and figure legend.

## Acknowledgments

We would like to thank Eric Peters for the assistance on mass spectrometry, Thomas Jentsch and Linyin Li for providing *LRRC8⁻/⁻* HEK293 cells and SWELL1 KO TIME cells, respectively. All cryo-EM data were collected at the Beckman Center for Cryo-EM at Johns Hopkins University. This work was supported by grants from the National Institute of Health (NIH R35GM124824, R01NS118014, and RF1NS134549 to Z.Q. and R35GM154904 to E.C.T.). Z.Q. was also supported by the American Heart Association Established Investigator Award, McKnight Scholar Award, Klingenstein-Simon Scholar Award, Sloan Research Fellowship in Neuroscience, and Randall Reed Scholar Award. E.C.T. was supported by the Searle Scholars Program (Kinship Foundation grant no. 22098168) and the Diana Helis Henry Medical Research Foundation (grant no. 142548).

## Author contributions

W.Z. performed molecular biology, biochemistry, and cGAMP assays. T.H. carried out protein purification and determined the cryo-EM structure. Hao W. conducted the majority of electrophysiological recordings. H.Y.C. performed cell sorting. N.K. and S.M. performed electrophysiological recordings of Jurkat and BV2 cells, respectively. K.H.C. performed live cell imaging. Haobo W. and A.K.M. helped with the cryo-EM study. W.Z., T.H., Hao W., E.C.T., Z.Q. analyzed the data. Z.Q. performed the proteomics experiments and supervised the project with E.C.T. Z.Q., W.Z., T.H. wrote the manuscript with input from all authors.

## Competing interests

The authors declare that they have no competing interests.

## Data availability

The atomic coordinates have been deposited in the Protein Data Bank, under the accession number 9O5K. Cryo-EM density maps have been deposited in the Electron Microscopy Data Bank (EMDB) under the accession number EMD-70143.

## Notes

### Competing Interest Statement

The authors have declared no competing interest.

